# dsRNA-induced condensation of antiviral proteins promotes PKR activation

**DOI:** 10.1101/2022.01.14.476399

**Authors:** Giulia A. Corbet, James M. Burke, Gaia R. Bublitz, Roy Parker

**Affiliations:** Department of Biochemistry, University of Colorado, Boulder 80303; Howard Hughes Medical Institute, Chevy Chase, MD 20815-6789

## Abstract

Mammalian cells respond to dsRNA in multiple manners. One key response to dsRNA is the activation of PKR, an eIF2α kinase, which triggers translational arrest and the formation of stress granules. However, the process of PKR activation in cells is not fully understood. In response to increased endogenous or exogenous dsRNA, we observed that PKR forms novel cytosolic condensates, referred to as dsRNA-induced foci (dRIFs). dRIFs contain dsRNA, form in proportion to dsRNA, and are enhanced by longer dsRNAs. dRIFs also enrich several other dsRNA-binding proteins including ADAR1, Stau1, NLRP1, and PACT. Strikingly, dRIFs correlate with and form prior to translation repression by PKR and localize to regions of cells where PKR activation is initiated. We suggest that dRIF formation is a mechanism cells utilize to enhance the sensitivity of PKR activation in response to low levels of dsRNA, or to overcome viral inhibitors of PKR activation.

## INTRODUCTION

Mammalian cells initiate cell-autonomous innate immune responses upon detection of double-stranded RNA (dsRNA), triggering a signaling cascade (reviewed in Jensen and Thomsen, 2012). This signaling cascade is initiated by numerous dsRNA sensors in the cell, also known as pattern recognition receptors (PRRs) (reviewed in Takeuchi and Akira, 2010). One of these PRRs is Protein kinase R (PKR), which binds dsRNA via its N-terminal RNA-binding domains and forms homodimers (Meurs et al., 1990; Nanduri et al., 2000). PKR dimerization on dsRNA results in autophosphorylation, leading to full activation of PKR catalytic activity (Nanduri et al., 2000; Ung et al., 2001; Zhang et al., 2001). Activated p-PKR phosphorylates the eukaryotic translation initiation factor eIF2α on serine 51 (p-eIF2α), which inhibits canonical AUG-dependent translation initiation (De Benedetti and Baglioni, 1984; Wek et al., 2006). This process shuts off bulk translation to reduce viral gene expression while promoting expression of select host mRNA transcripts involved in the integrated stress response (Dalet et al., 2015).

The inhibition of translation by PKR results in the disassociation of most cellular mRNAs from ribosomes. A fraction of these non-translating mRNAs condenses into cytoplasmic ribonucleoprotein complexes (RNPs) termed stress granules, which are enriched with large RNAs and several RNA-binding proteins: G3BP1/2, TIA-1, UBAP2L, poly(A)-binding protein (PABP) (Jain et al., 2016; Khong et al., 2017).

Various reports have reported the recruitment of dsRNA sensors, including PKR, MDA-5, RIG-I, and OAS, to stress granules assembled in response to dsRNA, viral infection, G3BP1 overexpression, or oxidative stress (Cadena et al., 2021; Langereis et al., 2013; Manivannan et al., 2020; Onomoto et al., 2012; Reineke and Lloyd, 2014; Yoo et al., 2014). These reports propose that interactions between stress granule proteins modulate the activation of dsRNA sensors to regulate the dsRNA response. For example, PKR localization to stress granules was proposed to promote phosphorylation of eIF2α (Reineke and Lloyd, 2014), and RIG-I localization to stress granules was proposed to promote the RIG-I/MAVS/IRF3 signaling pathway (Manivannan et al., 2020; Onomoto et al., 2012; Yoo et al., 2014).

While stress granules are presumed to promote the antiviral response, we recently showed that activation of the OAS/RNase L antiviral pathway inhibits the assembly of canonical stress granules by degrading cellular RNAs (Burke et al., 2019, 2020). Moreover, RNase L-mediated RNA decay promotes the formation of stress granule-like RNP complexes termed RNase L-dependent bodies (RLBs), which contain many stress granule enriched RNA-binding proteins: G3BP1/2, caprin-1, and PABP, but are distinct in their biogenesis, morphology, and composition in comparison to stress granules.

Here, we sought to determine how the localization and function of PKR occurs relative to cytoplasmic RNP granules. Surprisingly, in contrast to previous studies, we did not observe that PKR localized to stress granules. Instead, we observed that PKR forms novel foci in in response to foreign or endogenous dsRNA that are distinct from both RLBs and stress granules. These foci, which we have termed dsRNA-induced foci (dRIFs), contain dsRNA and various dsRNA-binding proteins, including PKR, ADAR1, Stau1, DHX9, NLRP1, and PACT. We observed that dRIF assembly is independent of PKR, indicating that these are not PKR-driven condensates. These findings identify a new type of RNA-protein condensate that forms during the innate antiviral response and may modulate the activity of PKR. Moreover, these findings challenge the notion that stress granules promote antiviral signaling by concentrating PRRs such as PKR and ADAR1.

## RESULTS

### PKR forms distinct foci during dsRNA stress

Intracellular dsRNA triggers activation of the innate immune response, which results in widespread RNA degradation, translation shutoff, and the formation of stress granules (SGs) or RNase L-dependent bodies (RLBs) (Burke et al., 2019; Jensen and Thomsen, 2012; Koyama et al., 2008; Yoo et al., 2014). Various studies have reported the localization of dsRNA-binding proteins and innate immune proteins, including PKR, to SGs during viral infection or other types of dsRNA stress (Cadena et al., 2021; Manivannan et al., 2020; Onomoto et al., 2012; Yoo et al., 2014). To see if we could reproduce this result, we stained cells for PKR and the SG marker G3BP1 after transfection of the synthetic dsRNA poly(I:C) into wild-type (WT) A549 cells (Fig. 1A).

**Figure 1.**
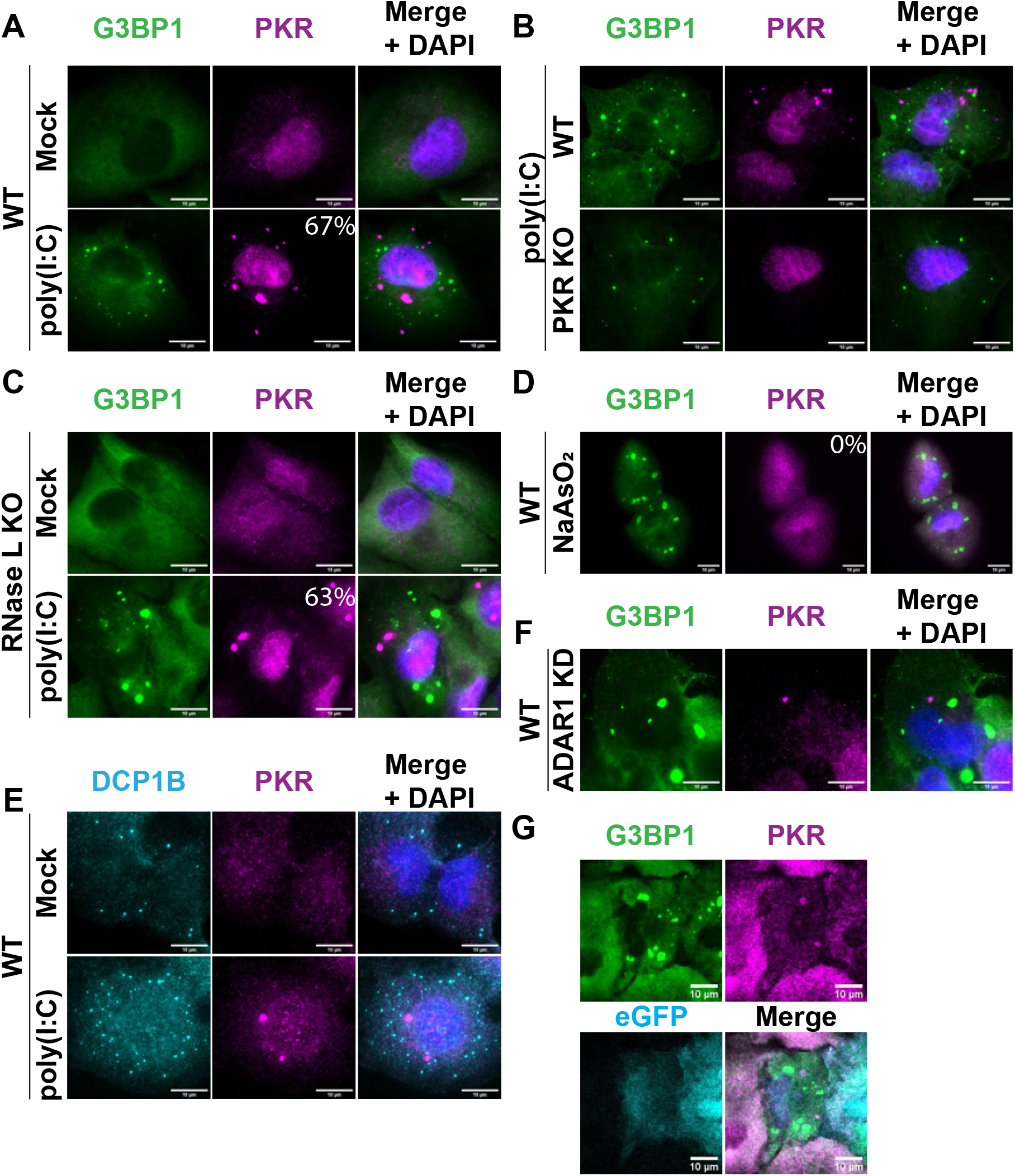
PKR forms distinct foci during dsRNA stress. A) Immunofluorescence (IF) for PKR and G3BP1 in wild-type (WT) A549 cells transfected with 500 ng/mL poly(I:C) for 4 hours. Nuclei stained with DAPI. Percentage of cells with PKR foci indicated. B) IF for PKR and G3BP1 in WT and PKR KO A549 cells transfected with poly(I:C). C) IF for PKR and G3BP1 in RNase L KO A549 cells transfected with poly(I:C). Percentage of cells with PKR foci indicated. D) IF for PKR and G3BP1 in A549 cells treated with 500 μM NaAsO_2_ for 1 hour. Percentage of cells with PKR foci indicated. E) IF for DCP1B and PKR in A549 cells transfected with poly(I:C). F) IF for PKR and G3BP1 in ADAR1 siRNA-treated U-2 OS cells. G) IF for PKR and G3BP1 in RL KO A549 cells transfected with eGFP-IRAlu reporter plasmid. All scale bars, 10 μm.

Surprisingly, we did not observe colocalization between PKR and G3BP1, and in fact, PKR formed distinct foci in 67% of cells (Fig. 1A). Similar results were observed in WT U-2 OS Cells (Fig. S1A). To validate that we were visualizing PKR, we performed the same poly(I:C) transfection and staining in PKR knockout (KO) A549 cells (Li et al., 2017). While some nonspecific antibody staining can be observed in the nuclei of PKR KO cells, we did not observe any PKR assemblies in the PKR KO cells (Fig. 1B), validating that the foci we observed in the WT cells contained PKR.

We could not directly rule out PKR localization to stress granules in the above experiment because WT A549 cells form small G3BP1 puncta known as RNase-L dependent bodies (RLBs) instead of stress granules (SGs) after poly(I:C) transfection because of widespread RNA degradation by RNase L (Fig. 1A) (Burke et al., 2019, 2020). To directly ask whether PKR would localize to SGs, we examined the subcellular localization of PKR and G3BP1 after poly(I:C) transfection into RNase L knockout (RL KO) A549 cells, which due to the absence of RNase L form SGs instead of RLBs (Burke et al., 2020). We observed that PKR formed distinct foci in the cytoplasm in 63% of cells and did not localize to SGs (Fig. 1C). Similar results were observed in RNase L KO U-2 OS Cells (Fig. S1B). We also did not observe PKR enrichment in SGs under arsenite stress by immunofluorescence or using an mApple-tagged PKR construct (Fig. 1D, S1C), suggesting that PKR does not localize to SGs in multiple different stresses. These observations demonstrate that PKR foci are distinct from SGs and RLBs.

One possibility is that PKR foci form due to PKR localizing upon poly(I:C) transfection to P-bodies, which are cytoplasmic RNP granules containing nontranslating mRNAs and the RNA decay machinery (Sheth and Parker, 2003). To test this possibility, we stained A549 cells for PKR and the P-body marker DCP1B. We did not observe colocalization between PKR foci and DCP1B-marked P-bodies (Fig. 1E). This demonstrates that PKR forms novel, distinct foci. Interestingly, in U-2 OS cells, we did observe that occasionally PKR assemblies are adjacent to P-bodies (Fig. S1D), suggesting there might be some interaction between PKR assemblies and P-bodies.

A recent report described that cytosolic dsRNA induces condensation of the inflammasome protein NLRP6 (Shen et al., 2021). To test whether the PKR assemblies we observed are related to NLRP6 condensates, we transfected a GFP-NLRP6 fusion protein into a cell line expressing mApple-PKR. Upon poly(I:C) transfection, we observed that mApple-PKR forms foci that do not recruit GFP-NLRP6 (Fig. S1E). Only in less than 15% of transfected cells did we observe the overexpressed GFP-NLRP6 transiently forming assemblies, and these also did not recruit mApple-PKR (Fig. S1E). These results indicate that PKR and NLRP6 form distinct assemblies in poly(I:C) stress, with PKR foci being more prevalent in A549 cells.

We examined whether formation of PKR foci was a common feature of dsRNA stress or whether it was restricted to exogenously transfected poly(I:C). We, and others, have previously observed that depletion of the dsRNA-modifying protein ADAR1 triggers PKR activation and stress granule assembly in a subset of cells (Berchtold et al., 2018; Corbet et al., 2021), presumably due to increased endogenous dsRNA from the lack of RNA editing. Thus, we tested whether ADAR1 KD induced the formation of PKR foci. We observed that in some WT U-2 OS cells, ADAR1 KD triggered the formation of PKR foci in the same cells that had stress granules, suggesting that PKR foci formation is linked to PKR activation (Fig. 1F). This observation demonstrates that endogenous dsRNAs can trigger PKR foci formation.

In a second experiment to examine if endogenous dsRNAs can induce PKR foci formation, we employed the use of a reporter plasmid expressing eGFP with an inverted *Alu* repeat (*IRAlu*) in the 3’ UTR (Chen et al., 2008). When transcribed, the *IRAlu* repeat forms approximately 300 bases of contiguous dsRNA structure, which should bind and activate PKR. Upon transient transfection of the reporter plasmid, we observed that a subset of eGFP-expressing cells formed PKR foci and had stress granules (Fig. 1G). This provides a second observation that endogenous dsRNAs can induce PKR foci formation and linking the formation of these foci to PKR activation and translational repression.

Taken together, these results demonstrate that both endogenous and exogenous dsRNA induce the formation of PKR foci, which we refer to as <u>d</u>s<u>R</u>NA-<u>i</u>nduced-<u>f</u>oci (dRIFs) henceforth.

### dRIFs contain dsRNA

Given that the presence of endogenous or exogenous dsRNA triggers the formation of dRIFs, we transfected cells with fluorescently labeled poly(I:C) and asked whether it colocalized with PKR (Fig. 2A). We observed that the poly(I:C) did colocalize with PKR. We further confirmed this colocalization of poly(I:C) with PKR by staining for dsRNA with the dsRNA specific K1 antibody (Fig. 2B). Additionally, live cell imaging of a cell line expressing mApple-PKR transfected with fluorescently labeled poly(I:C) demonstrated recruitment of PKR to dsRNA foci upon poly(I:C) entering the cell (Fig. S1F). This suggests a model where poly(I:C) functions as a scaffold for the formation of higher order PKR assemblies.

**Figure 2.**
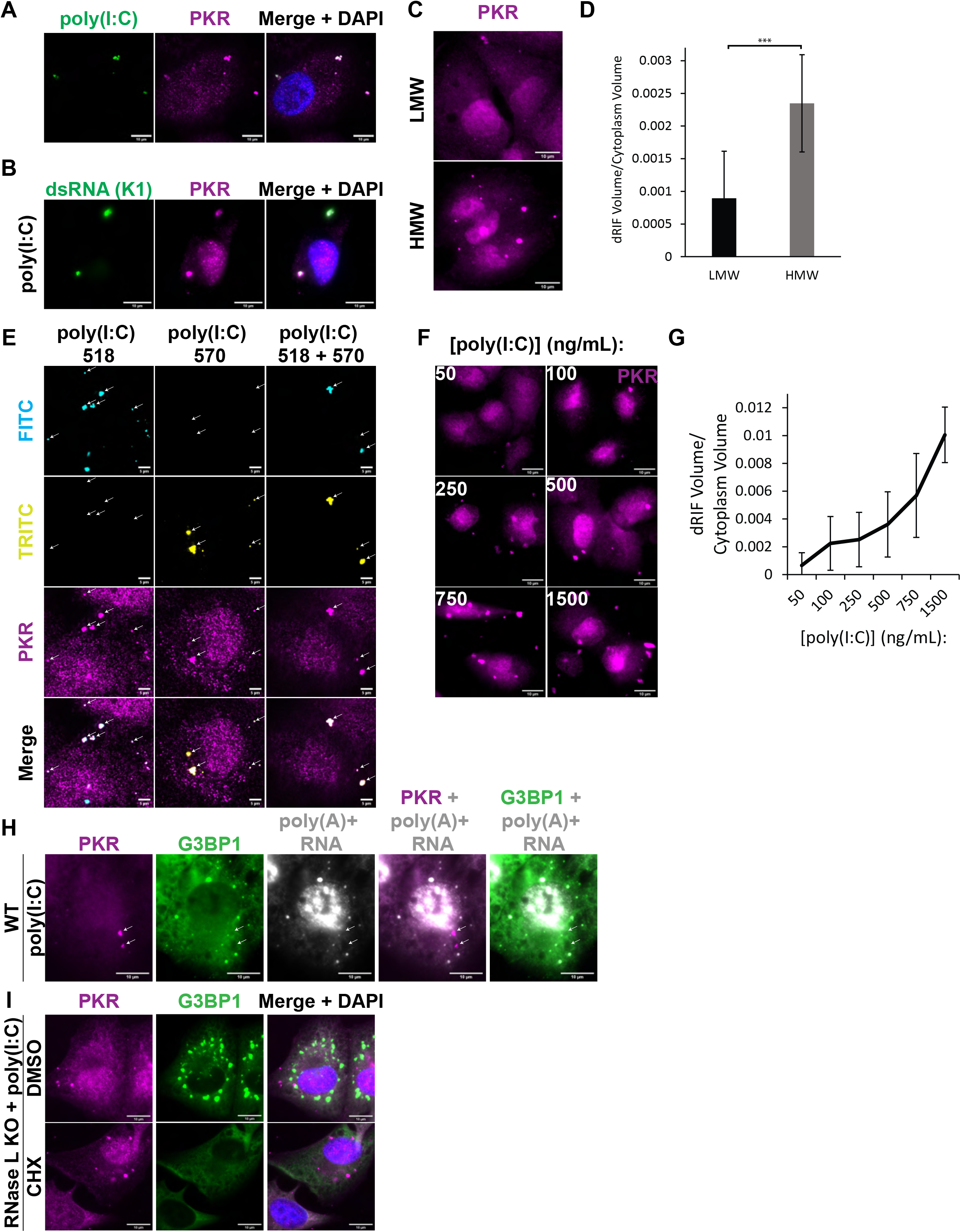
dRIFs contain dsRNA. A) IF for PKR in poly(I:C)-518-transfected A549 cells. Nuclei stained with DAPI. Scale bars, 10 μm. B) IF for PKR and dsRNA (K1 antibody) in A549 cells transfected with poly(I:C). Scale bars, 10 μm. C) IF for PKR in A549 cells transfected with 500 ng/mL of low molecular weight (LMW) or high molecular weight (HMW) poly(I:C) for 4 hours. Scale bars, 10 μm. D) Quantification of dRIF volume in C. 12 images from 2 independent experiments quantified. Error bars represent standard deviation. Unpaired two-tailed t-test, *** p < 0.001. E) IF for PKR in poly(I:C)-518 and poly(I:C)-570-transfected A549 cells. White arrows indicate location of dRIFs. Scale bars, 5 μm. F) IF for PKR in A549 cells transfected with 50-1500 ng/mL poly(I:C) for 4 hours. Scale bars, 10 μm. G) Quantification of dRIF volume relative to cytoplasmic volume in F. 5 images quantified for each condition. The line represents the average and error bars represent the standard deviation of dRIF volume measured at indicated concentrations of poly(I:C). H) IF for PKR and G3BP1, FISH for poly(A)+ RNA in A549 cells transfected with poly(I:C). White arrows indicate location of dRIFs. Scale bars, 10 μm. I) IF for PKR and G3BP1 in RL KO A549 cells treated with DMSO or 10 ug/mL cycloheximide (CHX) and transfected with 500 ng/mL poly(I:C) for 2 hours. Scale bars, 10 μm.

If poly(I:C) is the scaffold for dRIF formation, then the length of poly(I:C) would be expected to influence the size and/or number of dRIFs observed, with longer dsRNAs being more prone to forming dRIFs due to increased valency for interactions. To address this question, we transfected cells with equal nanograms of low molecular weight (LMW) and high molecular weight (HMW) poly(I:C). Because equal quantities of poly(I:C) were used, any differences in foci size and/or number can be explained by the difference in length of the poly(I:C) molecules.

We observed that high molecular weight poly(I:C) produces a larger volume of PKR foci than low molecular weight poly(I:C) for the same mass of transfected dsRNA (Fig. 2C, D). This result suggests that poly(I:C) serves as a scaffold for dRIF formation and that longer poly(I:C), which presumably contains more binding sites, can seed larger foci formation. Similar results have been seen with increased interaction sites enhancing protein-based condensate formation (Banani et al., 2016), and are consistent with the observations that longer RNAs are more effective at enhancing RNA condensation in vitro (Van Treeck et al., 2018), and in cells (Khong et al., 2017). Interestingly, no difference was observed in the phosphorylation of eIF2α upon transfection with equal doses of HMW or LMW poly(I:C), indicating that larger foci formation does not necessarily result in differences in PKR activation (Fig. S1G).

In principle, these assemblies could be composed of one or multiple dsRNA molecules bound to many PKR molecules. To test whether multiple dsRNA molecules are present in these foci, we transfected A549 cells with two different colors of fluorescently labeled poly(I:C). If these assemblies are composed of only one molecule of dsRNA, we should only ever observe one color of poly(I:C) in a single assembly. However, we observed assemblies that contained both colors of poly(I:C) (Fig. 2E), demonstrating that multiple molecules of dsRNA are present within each dRIF.

We wanted to test whether altering amounts of poly(I:C) had any effect on the formation of dRIFs. We transfected A549 cells with 50-1500 ng/mL poly(I:C) and stained for PKR. We observed a dose-dependent effect on dRIF volume with increasing amounts of poly(I:C) (Fig. 2F, G). This demonstrates a dose-response curve for dRIF formation dependent on the dsRNA concentration.

To determine if other RNAs could be recruited to dRIFs, we examined whether poly(A)+ RNA, which is enriched in SGs and RLBs, is also enriched in dRIFs. Upon staining for poly(A)+ RNA, the SG/RLB marker G3BP1, and PKR, we observed that poly(A)+ RNA is not enriched in dRIFs, but is enriched in RLBs, which stain positive for G3BP1 (Fig. 2H) (Burke et al., 2020). Together, these results demonstrate that the RNA content of dRIFs is primarily dsRNA.

Cycloheximide (CHX) is a translation inhibitor that traps mRNAs on ribosomes (Ennis and Lubin, 1964). mRNAs must be released from ribosomes to be recruited to SGs, thus, SG formation is inhibited by CHX (Kedersha et al., 2000; Mollet et al., 2008). Given that dRIFs are not enriched in poly(A)+ RNA, we would not expect dRIF formation to require mRNA release from ribosomes. To test this, we added 10 μg/mL CHX to RL KO A549 cells and transfected with 500 ng/mL poly(I:C) at the same time. After two hours, dRIF and SG formation was assessed by IF for PKR and G3BP1. While CHX prevents the formation of poly(I:C)-induced SGs, PKR recruitment to foci was not affected by CHX treatment (Fig. 2I), further confirming that non-translating mRNAs are not a major constituent of dRIFs.

Together, these observations demonstrate that dRIFs are assemblies of PKR and dsRNA. Both the average length and quantity of dsRNA added alter the volume of dRIFs formed, suggesting that dsRNA acts as a scaffold for dRIF assembly.

### dRIFs contain PKR-interacting proteins and dsRNA-binding proteins

Given that dRIFs appear to be composed of dsRNA and at least one dsRNA-binding protein, PKR, we hypothesized that other dsRNA-binding proteins might also enrich in dRIFs. The most immediately obvious set of proteins that appeared likely to enrich in dRIFs were dsRNA-binding proteins and PKR-interacting proteins. Thus, we performed IF for candidate proteins and assessed their localization upon poly(I:C) treatment.

Upon staining for the dsRNA-binding and modifying enzyme ADAR1, we observed frequent enrichment of ADAR1 in PKR foci upon poly(I:C) treatment (Fig. 3A). Approximately 85% of PKR+ foci were enriched for ADAR1 (Fig. 3A). Thus, ADAR1 is a component of dRIFs, but the ratios between ADAR1 and PKR can vary in dRIFs. ADAR1 recruitment to poly(I:C)-induced foci is unlikely to have functional consequences given that ADAR1 deaminates adenosines; however, ADAR1 recruitment to viral dsRNA-induced foci may have pro-viral or anti-viral consequences.

**Figure 3.**
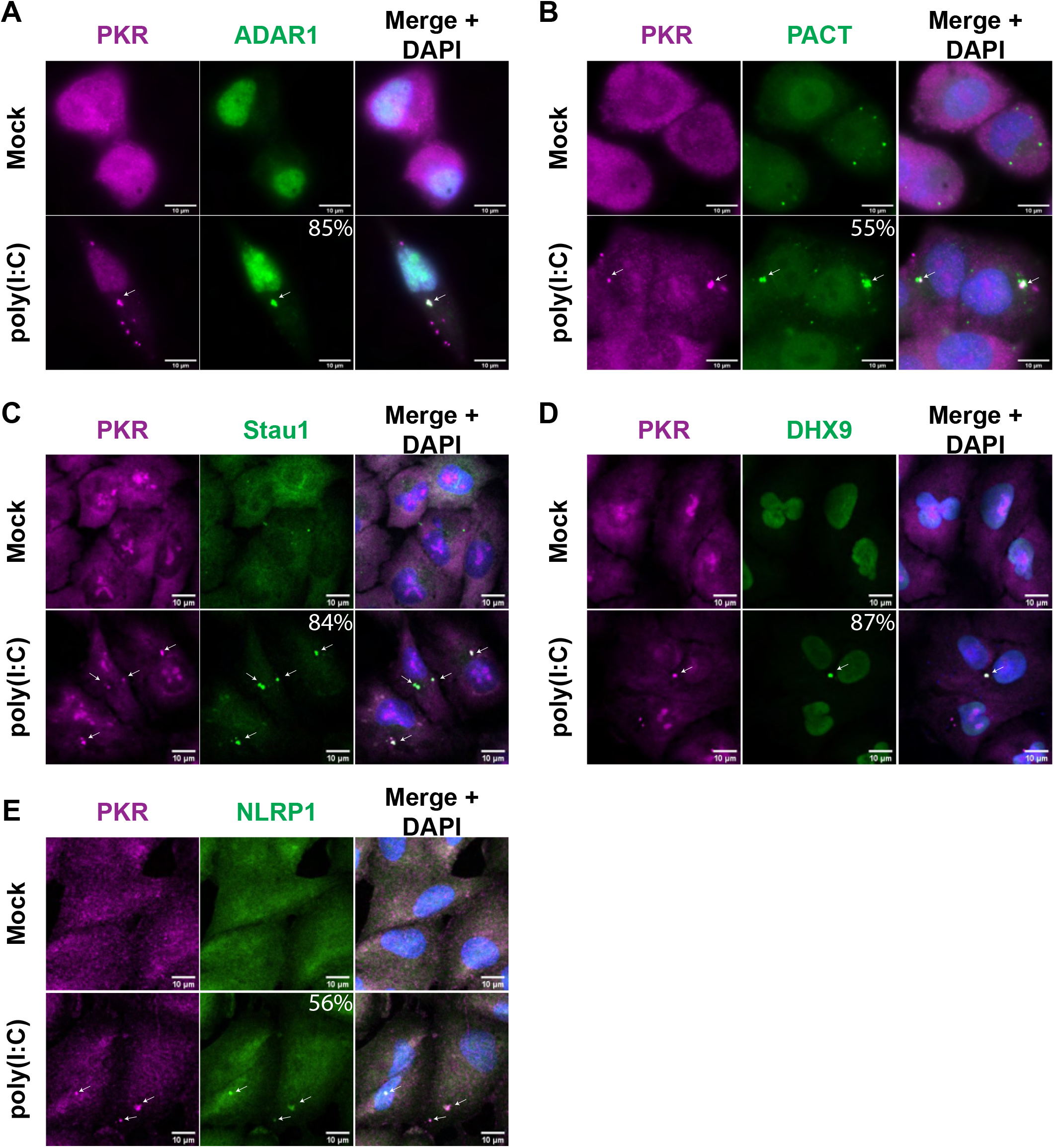
dRIFs contain dsRNA-binding proteins. A) IF for PKR and ADAR1 in A549 cells transfected with poly(I:C). White arrows indicate colocalization. Percentage of PKR+ dRIFs enriched for ADAR1 indicated. B) IF for PKR and PACT in A549 cells transfected with poly(I:C). Percentage of PKR+ dRIFs enriched for PACT indicated. C) IF for PKR and Staufen (Stau1) in A549 cells transfected with poly(I:C). Percentage of PKR+ dRIFs enriched for Stau11 indicated. D) IF for PKR and DHX9 in A549 cells transfected with poly(I:C). Percentage of PKR+ dRIFs enriched for DHX9 indicated. E) IF for PKR and NLRP1 in A549 cells transfected with poly(I:C). Percentage of PKR+ dRIFs enriched for NLRP1 indicated. All scale bars, 10 μm.

Protein activator of PKR (PACT) is another dsRNA-binding protein that is activated upon stress and activates PKR by direct binding (Farabaugh et al., 2020; Singh et al., 2011). Staining for PACT revealed that it is punctate even in non-stressed conditions (Fig. 3B). Upon poly(I:C) treatment, the PACT foci persisted and PACT was enriched in 55% of PKR+ foci (Fig. 3B). Live-cell imaging of a cell line expressing mApple-PACT and PKR-GFP validated the recruitment of both PKR and PACT to foci upon poly(I:C) transfection (Fig. S1H). While mApple-PACT does not form foci in the absence of added dsRNA, unlike what was observed when staining for endogenous PACT (Fig. 3B), it is recruited to foci upon poly(I:C) transfection.

Double-stranded RNA-binding protein Staufen homolog 1 (Stau1) is a highly conserved dsRNA-binding protein with roles in RNA localization, stability, and translation (Heraud-Farlow and Kiebler, 2014; Marión et al., 1999; Thomas et al., 2009). IF for Stau1 under mock conditions revealed it is largely diffuse in normal conditions, with a few small puncta detected (Fig. 3C). Upon poly(I:C) transfection, Stau1 is enriched in 84% of PKR+ dRIFs (Fig. 3C).

DHX9, also known as RNA Helicase A, is an essential dsRNA-binding protein with roles in RNA processing (Aktaş et al., 2017; Lee et al., 1998) IF for DHX9 revealed that it is diffuse throughout the nucleus and cytoplasm during normal conditions, but upon poly(I:C) treatment, it is enriched in 87% of PKR+ dRIFs (Fig. 3D). The inflammasome protein NLRP1 was recently demonstrated to have dsRNA-binding activity (Bauernfried et al., 2021) and appeared to be enriched in 56% of PKR+ foci (Fig. 3E).

Taken together, these results identify dRIFs as minimally containing dsRNA, PKR, PACT, ADAR1, Stau1, NLRP1, and DHX9. However, the enrichment of each of these dsRNA binding proteins varies, suggesting there is variation in the composition of each dRIF. Moreover, there appears to be selectivity in the recruitment of dsRNA binding proteins to dRIFs since several other dsRNA binding proteins did not enrich in dRIFs including NLRP3, NLRP6, TLR3, RIG-I, and Tudor-SN (Fig. S1E, S2A), although we cannot rule out that these proteins are present in dRIFs at levels similar to the bulk cytosol.

### dsRNA-binding enhances protein recruitment to dRIFs

In principle, the recruitment of proteins to dRIFs could be dependent on dsRNA binding, as well as proteinprotein interactions. To test the role of dsRNA binding for dRIF protein recruitment, we examined whether ADAR1’s dsRNA binding ability is necessary and/or sufficient for recruitment to dRIFs using previously generated truncation variants of the cytoplasmic isoform of ADAR1 (ADAR1 p150) (Corbet et al., 2021). ADAR1 p150 is known to localize to SGs induced by arsenite by its N-terminal Z-domains (Corbet et al., 2021; Ng et al., 2013). Thus, we asked whether various forms of ADAR1 p150 localize to dRIFs or SGs after poly(I:C) transfection.

We observed that in cells that formed both SGs and dRIFs in response to poly(I:C), ADAR1 p150 was recruited to dRIFs (Fig. 4A), validating our IF showing ADAR1 enrichment in dRIFs. Strikingly, we observed that the localization of ADAR1 p150 lacking its three dsRNA-binding domains (ΔdsRBD) to dRIFs is greatly reduced, and it is enriched in SGs in the majority of instances upon poly(I:C) transfection (Fig. 4A). This suggests that ADAR1 p150’s recruitment to dRIFs is largely dependent on its dsRNA-binding activity. Consistent with this interpretation, a construct expressing only ADAR1 p150’s three dsRNA-binding domains (dsRBD) localizes to dRIFs upon poly(I:C) transfection (Fig. 4A). These results demonstrate that ADAR1 dsRNA-binding domains are both necessary and sufficient for efficient ADAR1 p150 recruitment to dRIFs, and in the absence of dsRNA-binding, ADAR1 p150 can be recruited to SGs through its Z-domains (Corbet et al., 2021; Ng et al., 2013).

**Figure 4.**
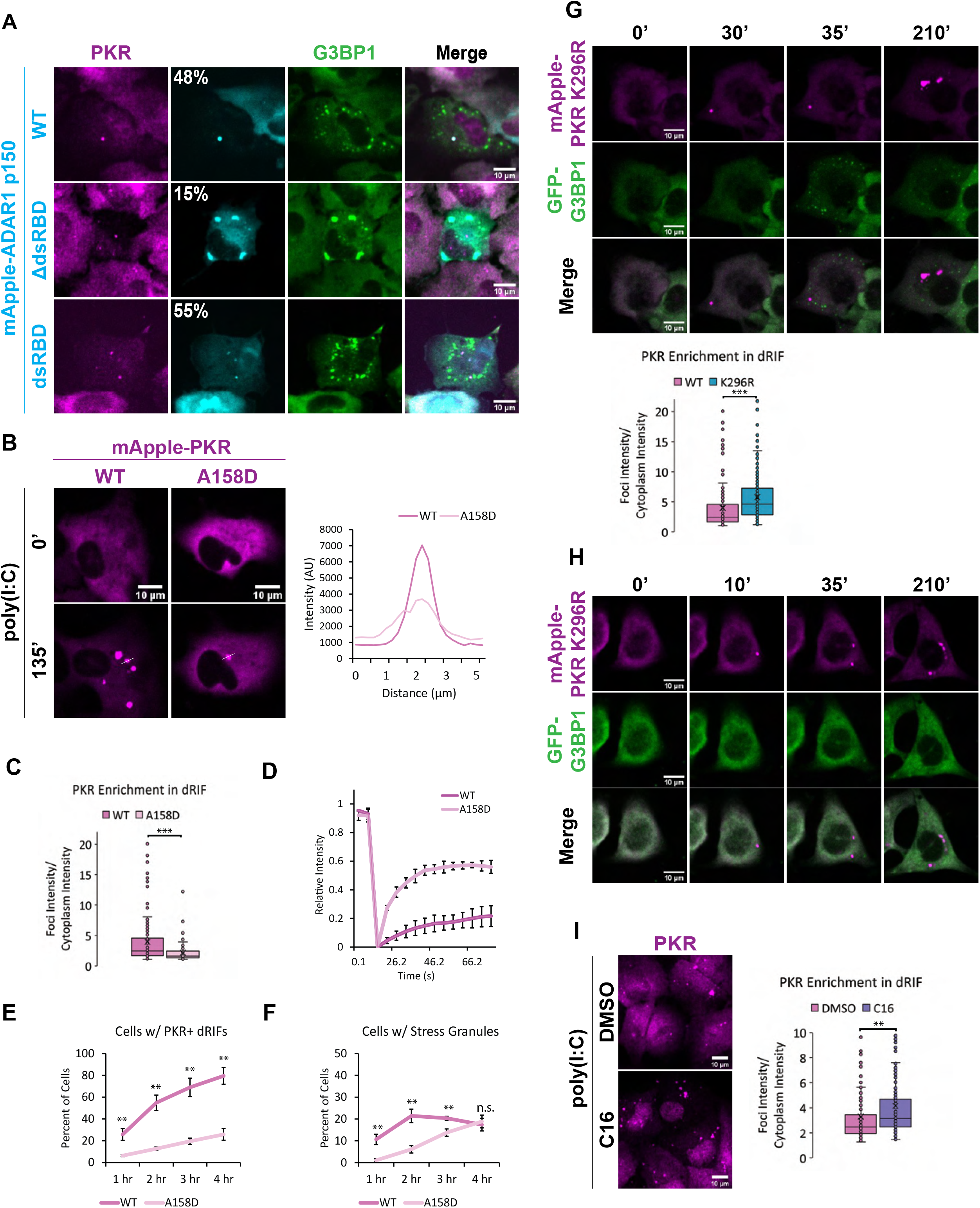
dsRNA-binding enhances protein recruitment to dRIFs. A) IF for PKR and G3BP1 in RL KO A549 cells transiently transfected with mApple-ADAR1 p150 WT, ΔdsRBD, or dsRBD and transfected with poly(I:C). Percentage of PKR+ dRIFs enriched for ADAR1 p150 indicated. B) A549 cells expressing mApple-PKR WT or A158D at 0- and 135-minutes after poly(I:C) transfection and a line scan showing the intensity profiles of foci. Scale bars, 10 μm. C) Quantification of average enrichment of mApple-PKR WT and A158D in dRIFs over cytoplasm. At least 83 foci quantified per condition from three independent experiments. Unpaired two-tailed t-test, *** p < 0.001. D) Fluorescence recovery after photobleaching (FRAP) analysis of mApple-PKR WT and A158D dRIFs. Error bars represent standard deviation. Data represents average from 3 independent experiments. E) Percentage of PKR/RL dKO A549 cells expressing GFP-G3BP1 and mApple-PKR WT or A158D with PKR+ dRIFs over time after poly(I:C) transfection. Data represents average from 3 independent experiments, error bars represent standard deviation. Unpaired two-tailed t-test, ** p < 0.01. F) Percentage of PKR/RL dKO A549 cells expressing GFP-G3BP1 and mApple-PKR WT or A158D with stress granules over time after poly(I:C) transfection. Data represents average from 3 independent experiments, error bars represent standard deviation. Unpaired two-tailed t-test, ** p < 0.01, n.s. = not significant. G) PKR KO A549 cells expressing GFP-G3BP1 and mApple-PKR K296R transfected with 500 ng/mL poly(I:C) and imaged at 0-, 30-, 35-, and 210-minutes after transfection. Quantification of mApple-PKR WT and K296R enrichment in dRIFs shown. At least 146 foci quantified from two or three independent experiments per condition. Unpaired two-tailed t-test, *** p < 0.001. H) PKR/RL dKO A549 cells expressing GFP-G3BP1 and mApple-PKR K296R transfected with 500 ng/mL poly(I:C) and imaged at 0-, 10-, 35-, and 210-minutes after transfection. I) IF for PKR in A549 cells treated with C16 PKR inhibitor or DMSO for 24 hours and transfected with 500 ng/mL poly(I:C) for 4 hours. Quantification of PKR enrichment in dRIFs in DMSO and C16-treated A549 cells. At least 169 foci quantified per condition from 4 independent experiments. Unpaired two-tailed t-test, ** p < 0.01. All scale bars, 10 μm.

To test the role of dsRNA binding in PKR’s recruitment to dRIFs, we utilized the A158D mutation which abolishes PKR’s ability to bind to dsRNA (Patel et al., 1996). Overexpression of mApple-PKR WT or A158D triggered stress granule formation in WT A549 cells, likely due to aberrant PKR activation and inhibition of translation initiation (Fig. S2B). This indicates that PKR is capable of auto-phosphorylation in the absence of dsRNA binding at high enough concentrations. Given this, we stably introduced mApple-PKR WT and A158D at close to endogenous levels using lentiviral transduction into PKR KO A549 cells.

We observed that mApple-PKR A158D is recruited to dRIFs upon poly(I:C) transfection; however, its recruitment is diminished compared to WT PKR (Fig. 4B, C, E). This observation suggests that PKR recruitment to dRIFs is a combination of binding dsRNA and protein-protein interactions between PKR and other dRIF proteins. Consistent with that model, FRAP analysis revealed that mApple-PKR A158D is much more mobile within dRIFs than WT, demonstrating that dsRNA-binding contributes to PKR’s lack of mobility within dRIFs (Fig. 4D). Live-cell imaging revealed that mApple-PKR WT is recruited to dRIFs quicker and in a higher proportion of cells than mApple-PKR A158D (Fig. 4E). This correlates to a higher proportion of mApple-PKR WT expressing cells having SGs, a readout of PKR-mediated translational shutoff, at earlier time points post-poly(I:C) transfection (Fig. 4F). By 4 hours post-transfection, no difference in the proportion of cells with SGs is observed between WT and A158D expressing cells (Fig. 4F). Since in vitro studies demonstrated that PKR A158D is unable to be activated by dsRNA (Patel and Sen, 1998), these results show that in cells, PKR A158D can still be activated by dsRNA, possibly due to its recruitment to and concentration in dRIFs via protein-protein interactions which facilitate PKR molecules being in close enough proximity to auto-phosphorylate.

One possible role of dRIFs is to concentrate PKR into a condensate with dsRNA and thereby increase the rate of PKR autophosphorylation *in trans*. Once phosphorylated, PKR’s affinity for dsRNA decreases, making PKR more likely to dissociate from dRIFs (V. Jammi and Beal, 2001). A prediction of this model is that inhibition of PKR catalysis by mutation or with chemical inhibitors should lead to prolonged persistence of PKR in dRIFs. To assess how PKR catalytic activity affects its recruitment to dRIFs, we generated a catalytically dead (K296R) mutant PKR and introduced it into PKR KO and PKR/RL dKO cells expressing GFP-G3BP1. Consistent with previous results (Burke et al., 2020), PKR KO cells expressing mApple-PKR K296R still form RLBs upon poly(I:C), because RNase L activation is not dependent on PKR activity (Fig. 4G). However, PKR/RL dKO cells expressing mApple-PKR K296R do not form SGs, because PKR-mediated phosphorylation of eIF2α is required for SG formation in response to poly(I:C) (Fig. 4H) (Burke et al., 2020). mApple-PKR K296R is more enriched in dRIFs than WT mApple-PKR, which is consistent with the non-phosphorylated form of PKR having a higher affinity for dsRNA (Fig. 4G).

C16 is a PKR inhibitor which binds to the ATP-binding pocket of PKR, preventing autophosphorylation, but does not prevent dsRNA-binding (Ingrand et al., 2007; Jammi et al., 2003). We tested whether inhibition of PKR with C16 would alter PKR recruitment to foci upon poly(I:C) treatment. After 24 hours of treatment with C16, A549 cells were transfected with poly(I:C) and stained for PKR. We observed an increase in PKR’s enrichment in dRIFs upon C16 treatment (Fig. 4I), which is again consistent with the unphosphorylated form of PKR having a higher affinity for dsRNA.

### dRIFs predominantly form prior to translational repression

To address the biological consequences of dRIF formation, we first examined the kinetics of dRIF formation relative to activation of PKR and RNase L. For these experiments, we generated a fluorescent mApple-tagged PKR construct. To avoid aberrant PKR activation due to overexpression, we stably introduced mApple-PKR and GFP-G3BP1 into a PKR KO A549 cell line using lentiviral transduction. PKR KO cells expressing mApple-PKR and GFP-G3BP1 did not form spontaneous stress granules in the vast majority of cells (Fig. 5A).

**Figure 5.**
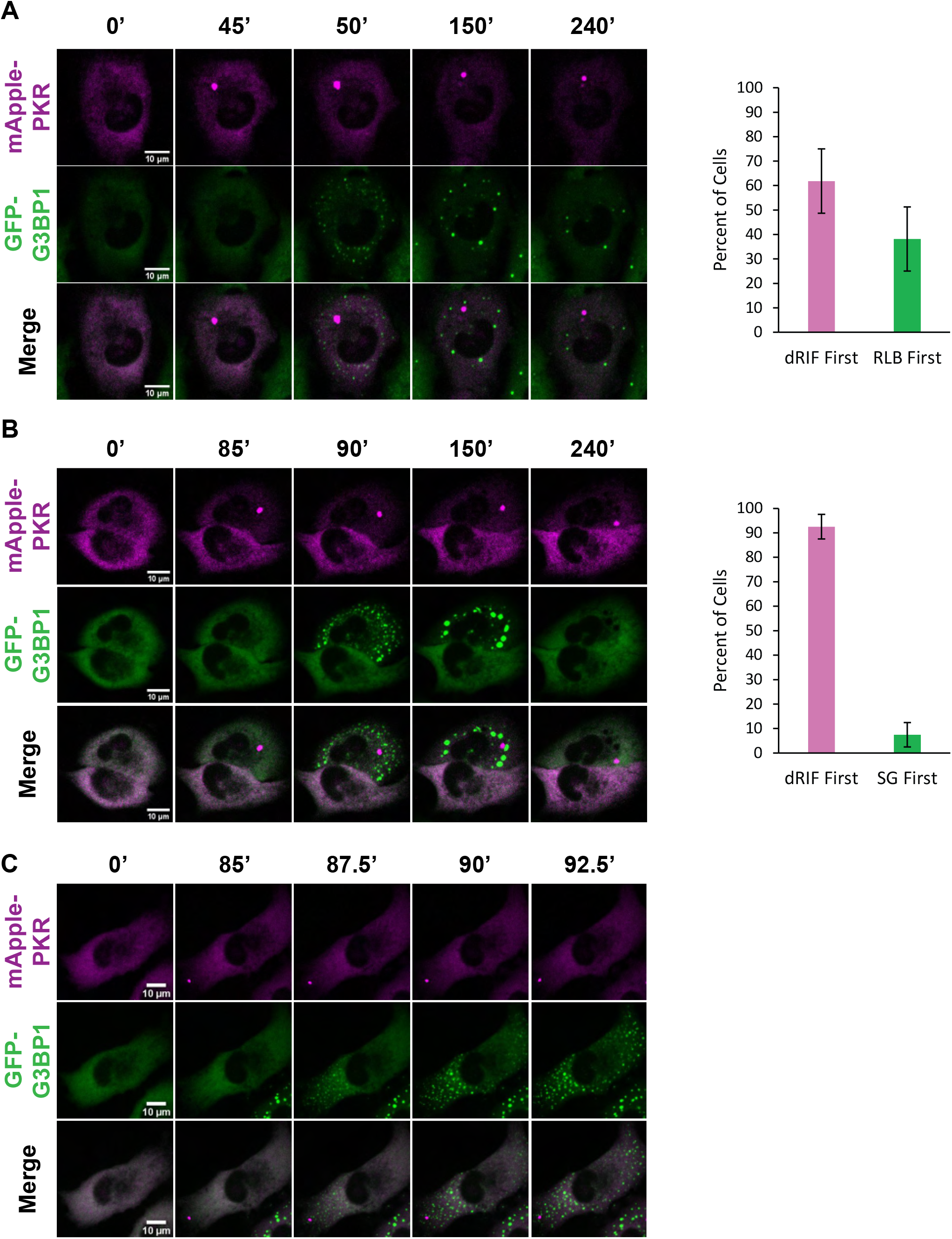
dRIFs predominantly form prior to translational repression. A) PKR KO A549 cells expressing GFP-G3BP1 and mApple-PKR transfected 500 ng/mL poly(I:C) and imaged at 0-, 45-, 50-, 150-, and 240-minutes after transfection. Percent of cells that form dRIFs first or RLBs first shown on the right. Error bars represent standard deviation from three independent experiments. B) PKR/RNase L double KO A549 cells expressing GFP-G3BP1 and mApple-PKR transfected with 500 ng/mL poly(I:C) and imaged at 0-, 85-, 90-, 150-, and 240-minutes after transfection. Percent of cells that form dRIFs first or SGs first shown on the right. Error bars represent standard deviation from three independent experiments. C) Images showing asymmetric SG formation proximal to dRIF in PKR/RNase L double KO A549 cells expressing GFP-G3BP1 and mApple-PKR transfected with 500 ng/mL poly(I:C) and imaged at 0-, 85-, 87.5-, 90-, and 92.5-minutes after transfection. All scale bars, 10 μm.

Simultaneous live-cell imaging of mApple-PKR, GFP-G3BP1 in PKR KO A549 cells transfected with poly(I:C) showed the formation of both dRIFs and RLBs, which are triggered by RNase L activation (Burke et al., 2020), typically within a few minutes of each other (Fig. 5A). In cells that formed both dRIFs and RLBs, dRIFs were observed forming before RLBs in 62% of cells, while RLBs were observed to form before dRIFs in 38% of cells (Fig. 5A). While the timing of RLB and dRIF formation varied from five minutes to several hours post-poly(I:C) transfection, cells that formed both types of assemblies typically formed both of them within 5-10 minutes of one another. This suggests that difference in the absolute time to a dsRNA response is likely due to variability in the timing of poly(I:C) release from liposomes. Moreover, these observations suggests that the timing of PKR foci formation and RNase L activation by poly(I:C) are similar, although they are independent from one another. Linescan analysis revealed that mApple-PKR was neither depleted nor enriched from RLBs, and GFP-G3BP1 was also neither depleted nor enriched in dRIFs (Fig. S2C). No instances of cells forming dRIFs, but not RLBs, were observed, validating that dsRNA is necessary for PKR recruitment to foci.

An important question is the timing of dRIF formation relative to PKR activation and translational repression, which can be assessed by the formation of stress granules in RNase L KO A549 cells. For this experiment, we stably introduced mApple-PKR and GFP-G3BP1 into PKR/RNase L double knockout (PKR/RL dKO) cells. Strikingly, in cells that formed both dRIFs and SGs upon poly(I:C) transfection, dRIFs formed prior to SGs in 93% of instances (Fig. 5B). This demonstrates that dRIFs typically form prior to PKR-mediated translation repression. Consistent with our fixed cell imaging, we observed that mApple-PKR and GFP-G3BP1 formed distinct assemblies upon poly(I:C) transfection, and neither was depleted nor enriched in the other assembly (Fig. 2B, S2D). In 7% of cells, SGs were observed prior to dRIFs, indicating that visible dRIF formation is not necessary for the formation of PKR-dependent SGs (Fig. S2E). However, we cannot rule out that smaller dRIF assemblies are forming that are below the detection limit of light microscopy, which can happen with RNP condensates (Rao and Parker, 2017).

In instances where dRIFs form near the edge of the cell, we can see that the first SGs form proximally to the dRIF, and then SG formation spreads across the cell (Fig. 5C, Video S1-5). This suggests that dRIFs represent sites of PKR activation and PKR phosphorylates the eIF2α molecules closest to the dRIF first, inhibiting translation locally before spreading across the cell. We interpret this observation to argue that dRIF formation is spatially linked to PKR activation and therefore dRIFs are sites of PKR activation.

RL KO cells will activate GADD34 to resume translation and disassemble SGs after poly(I:C) transfection (Burke et al., 2019, 2020; Clavarino et al., 2012). We observed that dRIFs persisted in cells when SGs were disassembled (Fig. 5B, Video S6). This observation argues that dRIF disassembly is not triggered by dephosphorylation of eIF2α or the resumption of active translation. We also observed that approximately one-third of cells formed dRIFs but did not form SGs during the duration of imaging. This shows that PKR recruitment to dRIFs does not always result in phosphorylation of eIF2α and translational repression (Fig. S2F).

These movies also reveal two properties of dRIFs similar to other biological condensates. First, demonstrating they share a dynamic consistent assembly mechanism, dRIFs can undergo both fusion and fission (Video S7, S8). Second, FRAP analysis of PKR shows that there are both dynamic and mobile pools of PKR in dRIFs, with the majority of PKR being relatively stably associated with dRIFs (Fig. 4F).

### PKR is not required for dRIF formation

An unresolved question is the mechanism by which dRIFs form. One possibility is that the multiple dsRNA binding domains of PKR form multivalent bridges between dsRNA molecules leading to dRIF condensation. To test this possibility, we stained WT and PKR KO A549 cells for dRIF proteins after poly(I:C) transfection. The localization of ADAR1, PACT, Stau1, and DHX9 to dRIFs were not affected by the loss of PKR (Fig. 6), demonstrating that dRIFs are not simply PKR-dependent assemblies. This suggests that the assembly of dRIFs is driven by other cellular factors, which remain to be identified.

**Figure 6.**
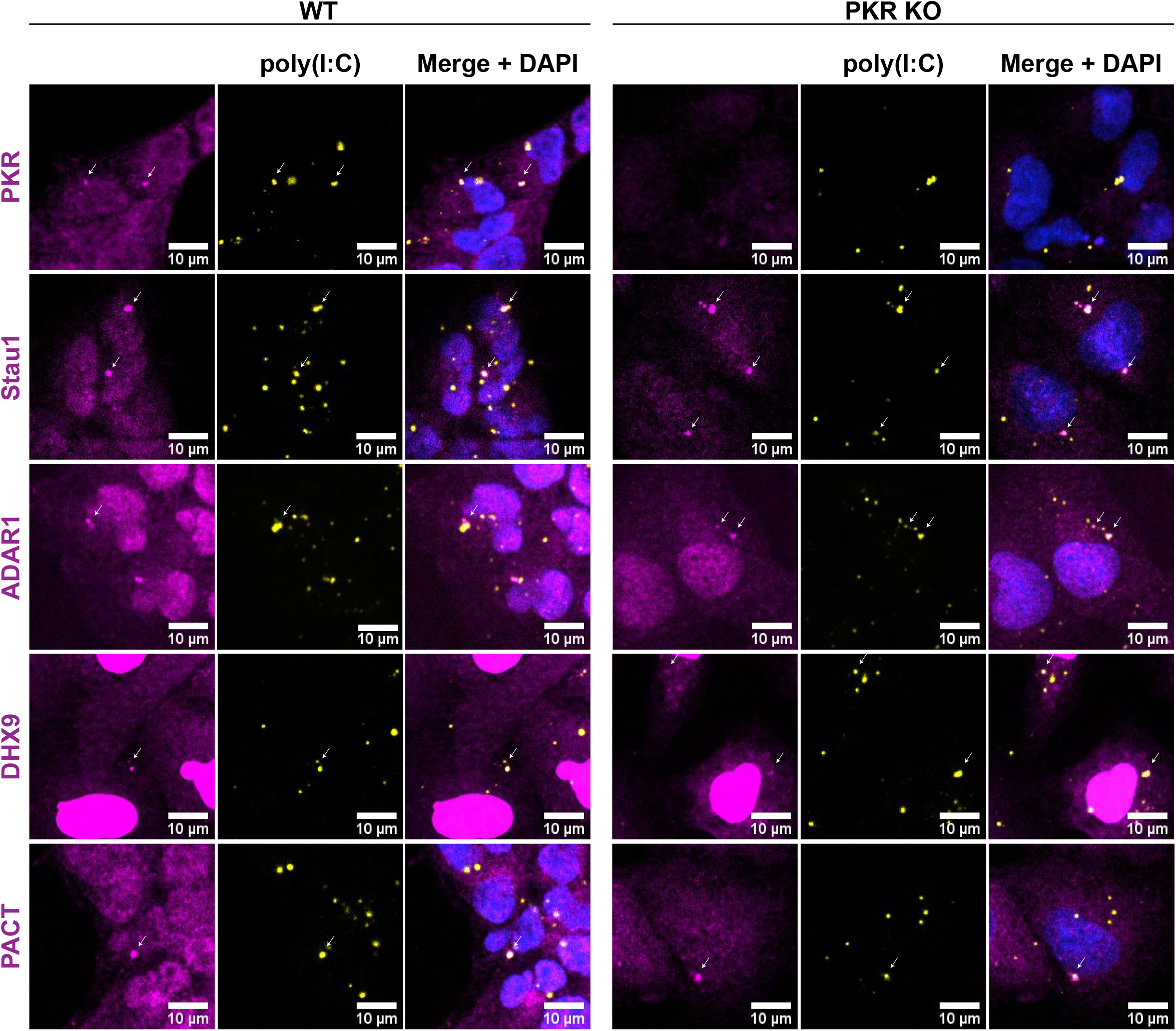
PKR is not required for dRIF formation. IF for PKR, Stau1, ADAR1, DHX9 and PACT in WT and PKR KO A549 cells transfected with labeled poly(I:C) for 4 hours. Nuclei shown in blue. White arrows indicate colocalization. All scale bars, 10 μm.

## DISCUSSION

In this work we present several observations documenting that cytosolic dsRNA can trigger the formation of a novel RNA and protein condensate in human cells referred to as <u>d</u>s<u>R</u>NA-<u>i</u>nduced <u>f</u>oci (dRIFs). First, we observe that dsRNA forms cytosolic foci following transfection (Figure 2). Second, those dsRNA foci recruit multiple, but not all, dsRNA-binding proteins, with PKR, ADAR1, PACT, Stau1, NLRP1, and DHX9 all showing increased partitioning into dRIFs (Figure 3). Moreover, dRIFs form in response to increased concentrations of endogenous dsRNA, either due to ADAR1 deficiency or expression of dsRNAs (Figure 1). dRIFs are distinct from other cytosolic RNP granules and do not overlap with stress granules, P-bodies, or RLBs (Figure 1). Consistent with these results, PKR foci have also been shown to form in response to dsRNA in another report (Zappa et al., 2021). Although dsRNA foci forming with NLRP6 have been previously described, dRIFs appear to be distinct since we do not observe any recruitment of NLRP6 to dRIFs (Figure S1).

Several observations argue that dRIFs are formed by dsRNA serving as a scaffold for proteins with multiple dsRNA binding domains, that can then introduce intermolecular protein-protein interactions between different dsRNA molecules (Figure 7). First, by using dsRNAs with different fluorescent tags, we demonstrate dRIFs contain multiple molecules of dsRNA (Figure 2). Second, longer dsRNAs are more efficient at generating dRIFs than shorter dsRNAs, even at the same mass of dsRNA (Figure 2). This is consistent with dRIF assembly being promoted by the increased valency of longer RNAs. Furthermore, many dsRNA binding proteins in dRIFs contain multiple dsRNA binding domains including Stau1, PACT, PKR, and ADAR1 (Figure 3). Given this, one possibility is that dRIF assembly is redundant with any of these proteins providing multivalent dsRNA binding to bridge dsRNA molecules and lead to dRIF formation.

**Figure 7.**
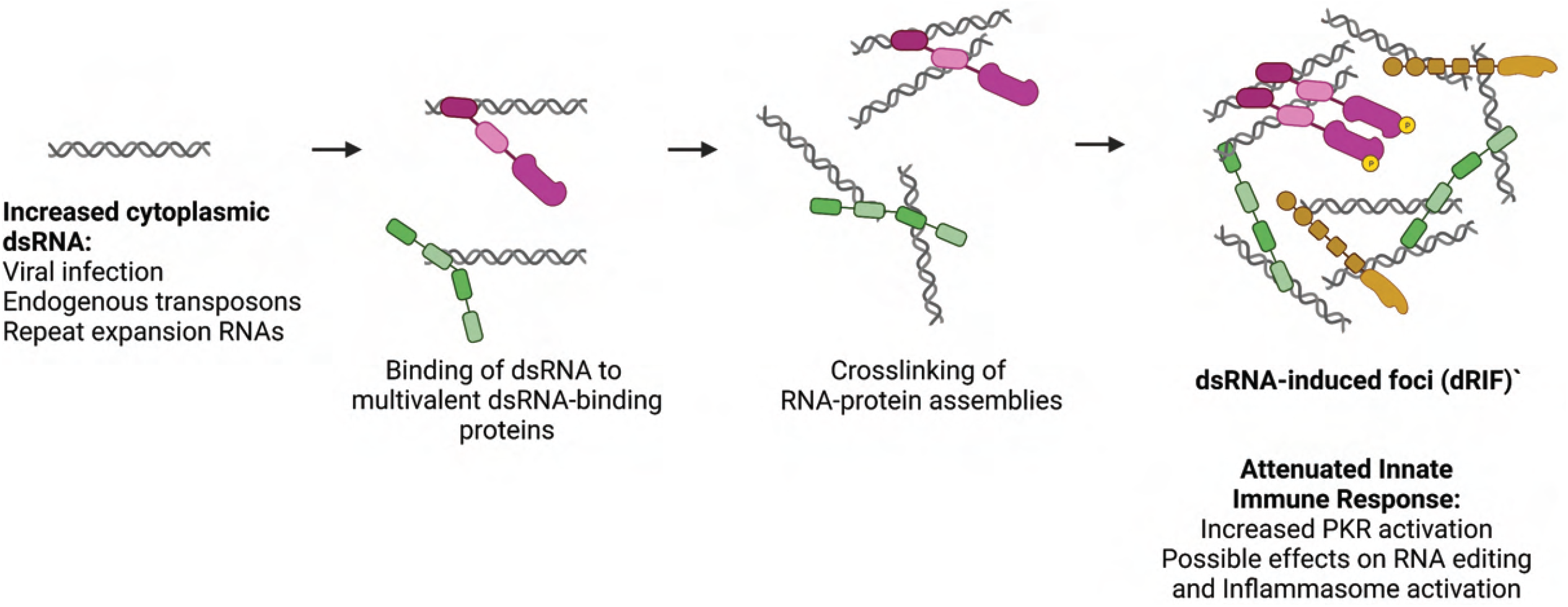
Model for dRIF formation. Under conditions of increased cytoplasmic dsRNA such as during viral infection, dsRNA-binding proteins bind to dsRNA. Multivalent protein-RNA and protein-protein interactions crosslink these RNA-protein assemblies, leading to the formation of a dsRNA-induced foci (dRIF), which results in increased PKR activation and may also have implications for RNA editing by ADAR1 and inflammasome activation by NLRP1. Created with BioRender.com

Our data provide several observations suggesting that dRIF formation will modulate the activation of PKR. First, we observe a correlation between cells that form visible dRIFs and those that trigger eIF2α phosphorylation and translation repression, as evidenced by stress granule formation (Figure 1F, 1G). Second, in cells lacking RNase L, which prevents RNase L-dependent eIF2α phosphorylation (Burke et al., 2019), we observe that dRIFs form before stress granules over 90% of the time (Figure 5). This argues dRIF formation precedes translation repression and may contribute to PKR activation. Third, in cells where dRIFs formation is asymmetric, we observe SGs first form close to the dRIF before spreading throughout the cell. This demonstrates a spatial correlation between dRIF formation and PKR activation, which is consistent with dRIFs serving as sites of PKR activation.

We anticipate that smaller dRIF assemblies form that we cannot observe in the light microscope and contribute to PKR activation for several reasons. In ~7% of RL KO cells we observed stress granules formed before dRIFs were observed, which could be due to smaller dRIFs forming, activating PKR, and then merging into a dRIF large enough to be visualized at a later time (Figure S2E).

The formation of dRIFs has implications for the activation of PKR in a number of biological contexts. We anticipate dRIFs will be important for PKR activation in some viral infections since viruses use multiple mechanisms to limit PKR activation (reviewed in Cesaro and Michiels, 2021). For example, hepatitis C virus blocks PKR dimerization through the NS5A protein (Gale et al., 1998, 1997). Given this, we hypothesize that the ability of cells to create a high local concentration of PKR in dRIFs will limit the ability of the NS5A protein to prevent PKR activation. Finally, we anticipate that dRIFs will be important in any biological context where the amount of dsRNA is limiting. These biological contexts include the initial phases of a viral infection, the higher levels of dsRNA in some Aicardi-Goutières syndromes (Rice et al., 2012), and even in neurological diseases where the expression of repeat expansion RNAs with dsRNA-like character can lead to activation of PKR (Peel et al., 2001; Tian et al., 2000). Consistent with a role for dRIFs in neurological disease, cells expressing the G4C2 repeat in the *C9orf72* gene that can cause ALS show activation of PKR and concentration of PKR into discreet cytoplasmic foci that we suggest are dRIFs (Rodriguez et al., 2021). Given these roles, an understanding of dRIF formation, the mechanisms by which dRIFs influence human disease, and how their manipulation may be therapeutic will be important areas of research.

The discovery of dRIFs adds to the growing set of observations whereby activation of the innate immune system through the recognition of either dsRNA or cytosolic DNA involves the formation of a nucleic acid-protein condensate. For example, the recognition of dsRNA by NLRP6 involves the formation of dsRNA-NLRP6 condensates (Shen et al., 2021), and our data suggests dRIFs may play a role in activating PKR and NLRP1 in response to dsRNA. Similarly, the recognition of cytosolic DNA by cGAS involves the formation of a DNA-protein condensate (Du and Chen, 2018). Thus, the formation of condensates that concentrate both the nucleic acid and the sensor provides a useful mechanism to increase the sensitivity of the innate immune response to nucleic acid.

## MATERIALS AND METHODS

### Plasmids

pGW1-mApple was a gift from Dr. Sami Barmada (Barmada et al., 2014). pGW1-mApple-PKR was generated from pGW1-mApple and pPET-PKR/PPase (Addgene Plasmid #42934) using In-Fusion® Snap Assembly (Takara Bio # 638947) and the following primers: GACGAGCTGTACAAGatggctggtgatctttcagca and GATCCGGTGGATCCCCTAACATGTGTGTCGTTCATTTTTC.

pGW1-mApple-PKR-K296R was generated from pGW1-mApple-PKR using Agilent QuikChange II XL Site-Directed Mutagenesis Kit (Neta Scientific # 200521) using the following primers: gttattatatttaacacgtctaataacgtaagtctttccgtcaattctgtgttttg and caaaacacagaattgacggaaagacttacgttattagacgtgttaaatataataac.

pLV-mApple-PKR WT and K296R were generated from pLV-EF1-Cre-PGK-Puro (Addgene Plasmid #108543) and pGW1-mApple-PKR using NEBuilder® HiFi DNA Assembly (NEB #E5520) and the following primers: gcaggctgccaccatgGTGAGCAAGGGCGAG and acaagaaagctgggtctaacatgtgtgtcg

pLV-mApple-PKR A158D was generated from pLV-mApple-PKR WT using Agilent QuikChange II XL Site-Directed Mutagenesis Kit and the following primers: tctgaagatatgcaagtttatcggccaattgttttgcttcc and ggaagcaaaacaattggccgataaacttgcatatcttcaga.

pPKR-GFP was generated from pGW1-mApple-PKR and pmGFP (Addgene # 117926) using NEBuilder® HiFi DNA Assembly with the following primers: ggagacccaagctggatggctggtgatctt and gcccttgctcaccatacatgtgtgtcgttc.

pLV-PKR-GFP was generated from pPKR-GFP and pLV-EF1-Cre-PGK-Puro using NEBuilder® HiFi DNA Assembly (NEB #E5520) and the following primers: gcaggctgccaccatggctggtgatctttca and acaagaaagctgggtctacttgtacagctc.

pGW1-mApple-PACT was generated from pcDNA-PACT (Addgene #15667) and pGW1-mApple using NEBuilder® HiFi DNA Assembly with the following primers: GACGAGCTGTACAAGATGTCCCAGAGCAGG and GATCCGGTGGATCCCCTACTTTCTTTCTGCTAT.

pmGFP-NLRP6 was generated from pmGFP and NLRP6 cDNA ORF Clone, Human, untagged (Sino Biological # HG21799-UT) using NEBuilder® HiFi DNA Assembly with the following primers: catggacgagctgtacaagATGGACCAGCCAGAG and gtttaaacgggccctctagaTCAGAAGGTCGAGATG.

pEGFP-IRAlus-Lin28 was a gift from Gordon Carmichael & Ling-Ling Chen (Addgene plasmid # 92347; http://n2t.net/addgene:92347; RRID:Addgene_92347).

pGW1-mApple-ADAR1p150 WT, ΔdsRBD, and dsRBD were generated as described in (Corbet et al., 2021).

### Cell Culture and Drug Treatment

The parental A549 cell line was provided by Dr. Christopher Sullivan (Burke et al., 2016). The PKR KO cell line was provided by Dr. Susan Weiss. The RNase L KO and PKR/RNase L double KO A549 and the RNase L KO U-2 OS cell lines were generated as described in (Burke et al., 2019). The WT U-2 OS cells were provided by Dr. Paul Anderson (Kedersha et al., 2008, 2016). All cell lines were maintained at 5% CO_2_ at 37 °C in Dulbecco’s Modified Eagle Medium (DMEM) supplemented with 10% fetal bovine serum (FBS) and 1% penicillin/streptomycin. Cells were routinely tested for mycoplasma contamination and were negative.

### siRNA Knockdowns and Plasmid Transfections

siRNA knockdowns were performed using Lipofectamine RNAiMAX (Thermo Fisher Cat # 13778075) according to Manufacturer’s protocol for 48 hours. The following siRNA was used: ADAR1 (Thermo Fisher Cat # 4390824).

Plasmids were transfected using jetPRIME® transfection reagent (VWR Scientific Cat # 89129-920) or Lipofectamine™ 2000 Transfection Reagent (Thermo Fisher Scientific #11668027) according to manufacturer’s protocol. Cell media was replaced 2 hours post-transfection. Cells were imaged or fixed 24 hours post transfection.

### Drug Treatments

Sodium arsenite (Millipore Sigma, S7400) was diluted in UltraPure DNase/RNase-Free Distilled Water (Thermo Fisher Scientific, 10977-023). Treatment was done for 1 hr at 37 °C. C16 (Imidazolo-oxindole PKR inhibitor C16,≥98% (HPLC), Sigma-Aldrich #I9785-5MG) was prepared in DMSO and treatments were done for 24 hours at 37 °C. Poly(I:C) (InvivoGen tlrl-pic, tlrl-picf, tlrl-picr, tlrl-picw) was resuspended in UltraPure DNase/RNase-Free Distilled Water and transfected into cells using Lipofectamine 2000 reagent (Thermo Fisher Scientific, 11668030) according to the manufacturer’s protocol. 3 μl Lipofectamine was used for 1 μg poly(I:C). Poly(I:C) treatments were done for 4 hours unless otherwise stated in figure legend. Cycloheximide (Sigma-Aldrich, C7698-1G) was prepared in DMSO, and treatment was performed for 2 hrs.

### Lentiviral Transductions

Lentivirus was generated in HEK293T cells. HEK293T cells in T-25 flask were transfected with 2 μg pLV-mApple-PKR (WT, A158D, or K296R), pLV-PKR-GFP, or pLenti–EF1–GFP–G3BP1–blast (Burke et al., 2020), 1 μg pRSV-Rev, 1 μg pVSV-G, and 1 μg of pMDLg–pRRE and 15 μL Lipofectamine 2000. Media was collected 48 hours post-transfection and filtered through a 0.45 μm filter. To generate stable cell lines, A549 cells were incubated 1 mL of lentivirus stock containing 10 μg/mL polybrene for 1 hour, at which point fresh media was added to the cells. 48 hours post-transduction, cells were selected with 1.5 μg/mL puromycin and/or blasticidin-containing media.

### Immunofluorescence

Immunofluorescence was performed as described in (Corbet et al., 2021). The following antibodies were used at 1:500 dilutions: G3BP1 antibody (Abcam ab56574), PABPC1 (abcam ab21060), ADAR1 (abcam ab88574), PACT (Abcam ab31967), PKR (Cell Signaling Technology #122976), PKR (Santa Cruz Biotechnology #sc-6282), Stau1 (Novus Biologicals #NBP1-33202), DHX9 (Proteintech C827D22), DCP1A (Millipore Sigma WH0055802M6-50UG), DCP1B (Cell Signaling Technology #13233S), TLR3 (Abcam ab137722), NLRP1 (Abcam ab36852), NLRP3 (Thermo Scientific PA5-109211). The following antibodies were used at 1:1000 dilutions: K1 (Cell Signaling Technology #28764S). The following secondary antibodies were used at 1:1000: Goat anti-rabbit (abcam ab6717), Goat anti-rabbit (abcam ab150079), Goat anti-mouse (abcam ab150115), Goat anti-mouse (abcam ab6785), Donkey anti-rabbit (abcam ab150062).

### Sequential Immunofluorescence and FISH

Sequential immunofluorescence and FISH was performed as described in (Khong et al., 2018). Poly d(T) probes were ordered from Integrated DNA Technologies as 30 deoxythymidines with a 5’ Cy5 modification.

### SDS-PAGE and Western Blotting

SDS-PAGE and Western blotting was performed as described in (Corbet et al., 2021). The p-eIF2α (Cell Signaling Technology #9721S) and eIF2α (Cell Signaling Technology #9722S) antibodies were used at 1:1000. The anti-rabbit IgG, HRP-linked antibody (Cell Signaling 7074S) was used at 1:2000.

### Microscopy and Image Analysis

A549 cells were grown in 24-well glass plates (Cellvis #P24-1.5H-N) or on coverslips (Thomas Scientific # 1217N79). Live cell imaging was performed on a Nikon Spinning Disc Confocal microscope with a 2x Andor Ultra 888 EMCCD camera and a 40x air objective. Fixed cells were imaged on a widefield DeltaVision Elite microscope with a 100X oil objective and PCO Edge sCMOS camera or the Nikon Spinning Disc Confocal microscope with a 2x Andor Ultra 888 EMCCD camera and a 40x air or 100x oil objective. 7-20 z slices were imaged at 0.2 μm (DeltaVision or Spinning Disc 100x) or 0.6 μm (Spinning Disc 40x) distance between slices. All images shown are of a single z plane unless otherwise specified in figure legend. Deconvolved images indicated in figure legend when shown.

dRIF enrichment was calculated using ImageJ. Maximum intensity projections of all z slices were generated, a line was drawn through the dRIF and through the cytoplasm, and the average intensity of each line was calculated. Enrichment was calculated as dRIF intensity/cytoplasm intensity. dRIF and cytoplasm volume were calculated using Bitplane Imaris software as described in (Corbet et al., 2021; Khong et al., 2017) for stress granule quantification. Statistical analysis was performed using an unpaired two-tailed t-test. For p > 0.05, analyses were designated as not significant.

### FRAP

A549 cells were grown in 24-well glass plates (Cellvis #P24-1.5H-N). Cell media was replaced with OptiMEM media immediately prior to imaging. FRAP was done on a Nikon A1R Laser Scanning Confocal microscope as follows: acquire 2 images 5 second apart prior to bleaching, bleach for 9.54 seconds with 405 and 488 nm lasers at 100% laser power, image every 5 seconds second for 1 minute during recovery. Mean intensity within the bleached area was determined using Nikon Elements software. Intensities were normalized to the average intensity of the region of interest (ROI) at time 0 (set to 1) and at the first time-point post-bleaching (set to 0). Graphs represent averages of three independent experiments where at least 10 foci were bleached for each experiment. Error bars represent standard deviation.

## ACKNOWLEDGEMENTS

We thank Dr. Nancy Kedersha and Dr. Paul Anderson for the parental, G3BP1/2-KO, and GFP–G3BP1 U-2 OS cell lines. We thank Dr. Susan Weiss for the A549 PKR KO cell line. The imaging work was performed at the BioFrontiers Institute Advanced Light Microscopy Core (RRID: SCR_018302). Spinning disc confocal microscopy was performed on Nikon Ti-E microscope supported by the BioFrontiers Institute and the Howard Hughes Medical Institute.

---

**Supplemental Figure 1.** A) IF for PKR and G3BP1 in WT U-2 OS cells transfected with poly(I:C) for 4 hours. Nuclei shown in blue. B) IF for PKR and G3BP1 in RL KO U-2 OS cells transfected with poly(I:C). C) PKR KO A549 cells expressing GFP-G3BP1 and mApple-PKR treated with 500 μM NaAsO_2_ for 45 minutes. D) IF for DCP1A and PKR in WT U-2 OS cells transfected with poly(I:C). E) PKR KO A549 cells expressing mApple-PKR and GFP-NLRP6 transfected with 500 ng/mL poly(I:C) and imaged 0-, 80-, 200-, and 360-minutes after transfection. F) PKR KO A549 cells expressing mApple-PKR and transfected with labeled poly(I:C) and imaged at 0-, 40-, 47.5-, and 50-minutes after transfection. G) Western Blot of p-eIF2α and eIF2α in RL KO A549 cells transfected with mock, 50 ng/mL, and 500 ng/mL high molecular weight (HMW) or low molecular weight (LMW) poly(I:C) for 4 hours. H) PKR KO A549 cells expressing mApple-PACT and PKR-GFP transfected with poly(I:C) and imaged at 0-, 85-, 160-, and 195-minutes after transfection. All scale bars, 10 μm.

**Supplemental Figure 2.** A) IF for PKR and dsRNA-binding proteins NLRP3, TLR3, RIG-I, and Tudor-SN in A549 cells transfected with poly(I:C) for 4 hours. Nuclei shown in blue. B) WT A549 cells transiently transfected with mApple-PKR WT or A158D and stained for G3BP1. C) Line-scan of PKR KO A549 cells expressing mApple-PKR and GFP-G3BP1 after poly(I:C) transfection. Maximum intensity in the line scan of each channel set to 1. D) Linescan of PKR/RNase L dKO A549 cells expressing mApple-PKR and GFP-G3BP1 after poly(I:C) transfection. Maximum intensity in the line scan of each channel set to 1. E) Images demonstrating SGs forming prior to dRIFs in PKR/RNase L dKO A549 cells expressing GFP-G3BP1 and mApple-PKR transfected 500 ng/mL poly(I:C) and imaged at 0-, 25-, 35-, and 220-minutes after transfection. F) Images demonstrating dRIFs forming without SGs in PKR/RL dKO A549 cells expressing GFP-G3BP1 and mApple-PKR transfected 500 ng/mL poly(I:C) and imaged at 0-, 105-, 150-, 220-minutes after transfection. All scale bars, 10 μm.

**Video S1-S5.** Video showing asymmetrical formation of GFP-G3BP1 stress granules (green) near site of mApple-PKR foci (magenta) and spreading across cell. Scale bar, 10 μm. Time since poly(I:C) transfection indicated.

**Video S6.** Video showing persistence of mApple-PKR foci (magenta) after dissolution of GFP-G3BP1 stress granules (green) formed after poly(I:C) transfection. Scale bar, 10 μm. Time since poly(I:C) transfection indicated.

**Video S7.** Video showing fusion of mApple-PKR foci formed after poly(I:C) transfection. Scale bar, 10 μm. Time since poly(I:C) transfection indicated.

**Video S8.** Video showing fission of mApple-PKR foci formed after poly(I:C) transfection. Scale bar, 10 μm. Time since poly(I:C) transfection indicated.

