## Supplementary figures and images for "dsRNA-induced condensation of antiviral proteins promotes PKR activation"

### Supplemental Figures

#### SUPPLEMENTAL FIGURE 1

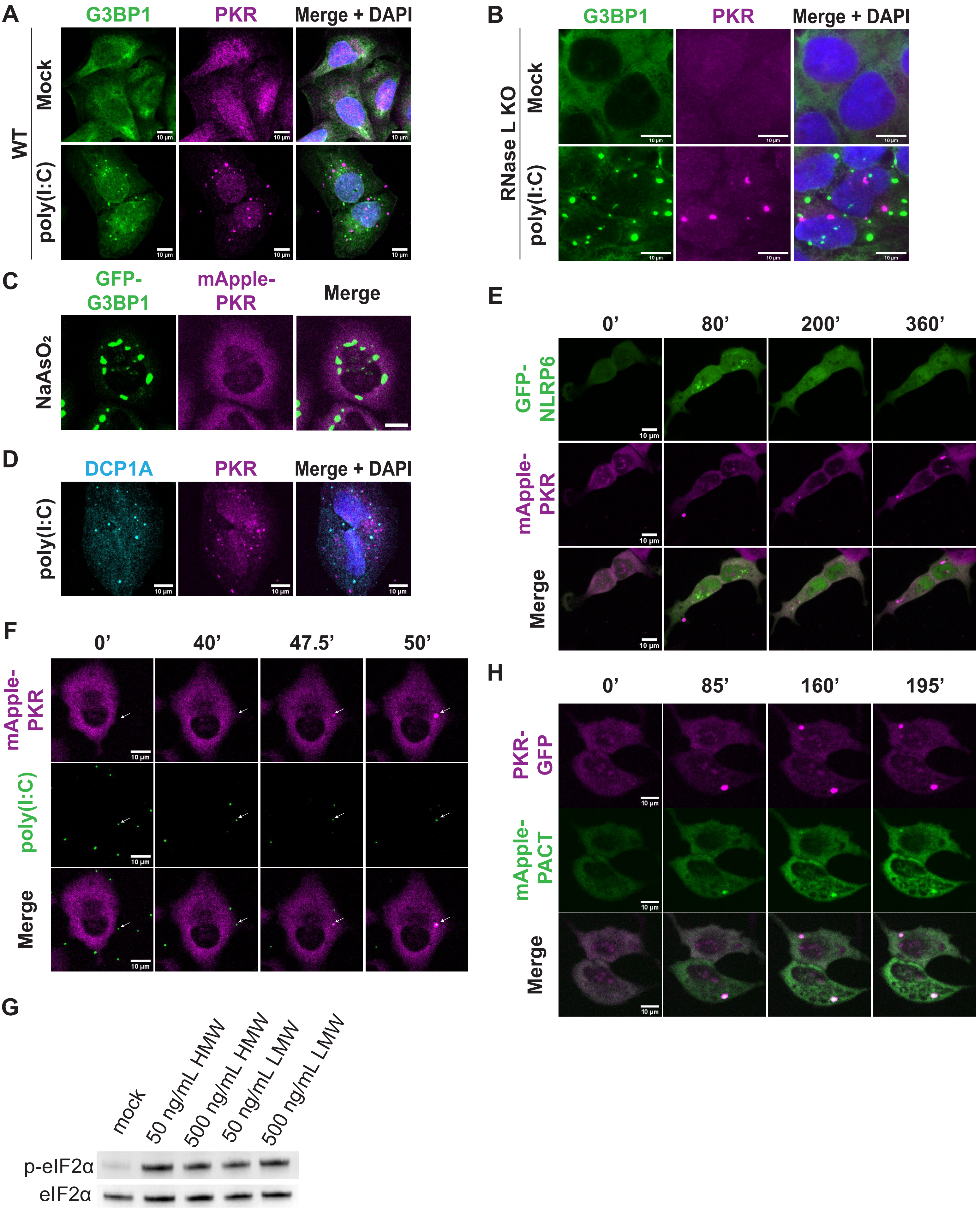

#### SUPPLEMENTAL FIGURE 2

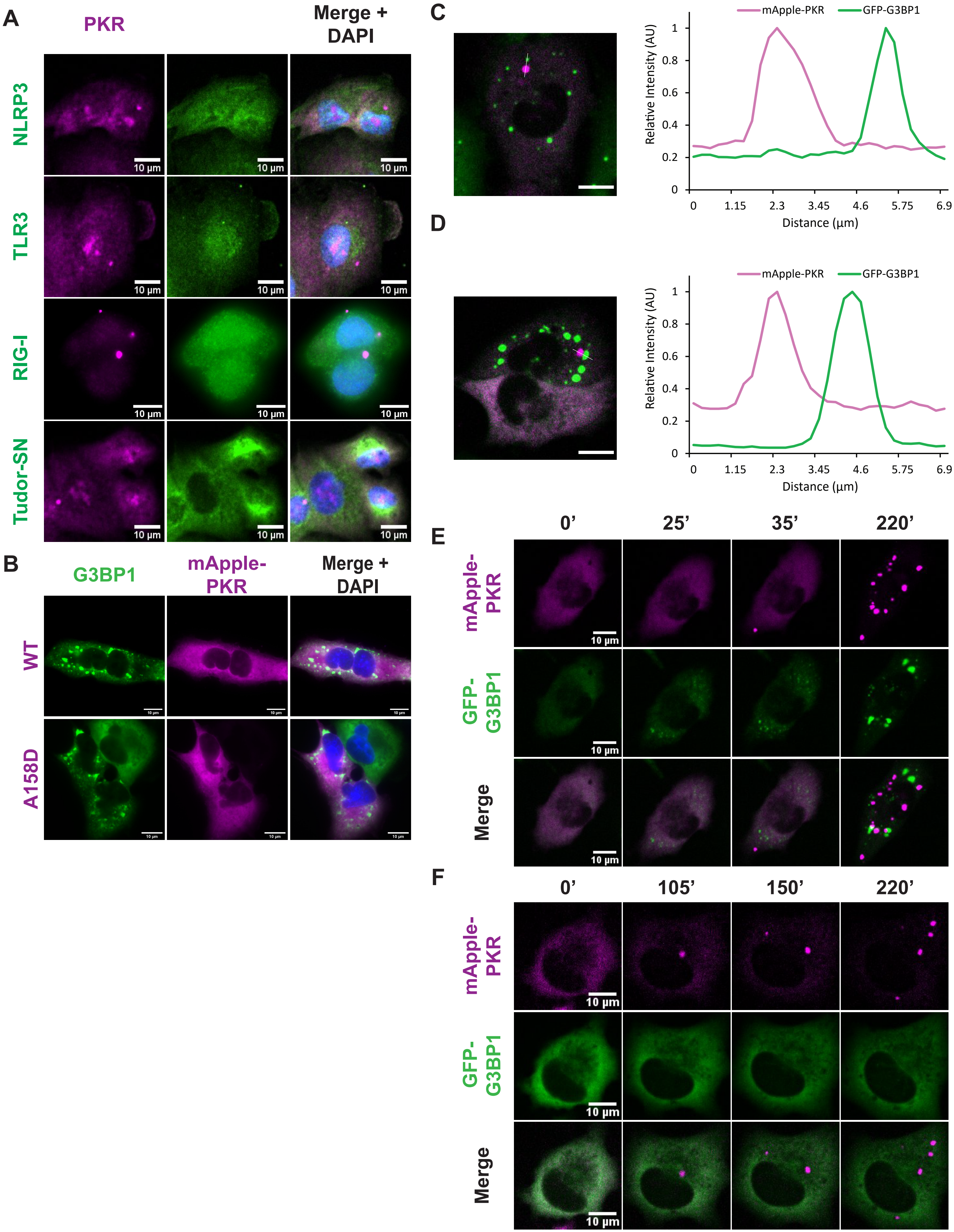
